# Concerning the eXclusion in human genomics: The choice of sex chromosome representation in the human genome drastically affects number of identified variants

**DOI:** 10.1101/2023.02.22.529542

**Authors:** Brendan J. Pinto, Brian O’Connor, Michael C. Schatz, Samantha Zarate, Melissa A. Wilson

## Abstract

Over the past 30 years, a community of scientists have pieced together every base pair of the human reference genome from telomere-to-telomere. Interestingly, most human genomics studies omit more than 5% of the genome from their analyses. Under ‘normal’ circumstances, omitting any chromosome(s) from analysis of the human genome would be reason for concern—the exception being the sex chromosomes. Sex chromosomes in eutherians share an evolutionary origin as an ancestral pair of autosomes. In humans, they share three regions of high sequence identity (~98-100%), which—along with the unique transmission patterns of the sex chromosomes—introduce technical artifacts into genomic analyses. However, the human X chromosome bears numerous important genes—including more “immune response” genes than any other chromosome—which makes its exclusion irresponsible when sex differences across human diseases are widespread. To better characterize the effect that including/excluding the X chromosome may have on variants called, we conducted a pilot study on the Terra cloud platform to replicate a subset of standard genomic practices using both the CHM13 reference genome and sex chromosome complement-aware (SCC-aware) reference genome. We compared quality of variant calling, expression quantification, and allele-specific expression using these two reference genome versions across 50 human samples from the Genotype-Tissue-Expression consortium annotated as females. We found that after correction, the whole X chromosome (100%) can generate reliable variant calls—allowing for the inclusion of the whole genome in human genomics analyses as a departure from the status quo of omitting the sex chromosomes from empirical and clinical genomics studies.

## Background

The X and Y chromosomes in placental mammals share an evolutionary origin as an ancestral pair of autosomes (Graves, 2008). Due to this shared ancestry and subsequent chromosomal rearrangements, the X and Y chromosomes in humans are highly divergent yet share regions of high sequence identity (~98-100%; Figure 1a), which introduces regions of varying ploidy across this chromosomal pair. Although this is well understood biologically, it introduces technical artifacts within modern genomic analyses that require correction to prevent potentially erroneous conclusions (Carey et al. 2022; Webster et al. 2019). Though these technical artifacts have remained ignored in many empirical and clinical studies, they have been used as justification to ignore the sex chromosomes on a grand scale and, therefore, the importance of sex-linked variation to human health is likely greatly underestimated (Inkster et al. 2023; Khramtsova et al. 2019; Köferle et al. 2022; Natri et al. 2019; Sun et al. 2023; Wise et al. 2013). Here, we aim to better grasp the scope of data lost by excluding or misrepresenting the sex chromosomes in human genomics. We urge empiricists and clinicians to confront these issues moving forward to simultaneously increase the number of genome-wide association studies (GWAS) and reduce the numbers of autosome-wide association scan studies (AWAS) currently being published (Sun et al. 2023).

**Figure 1:**
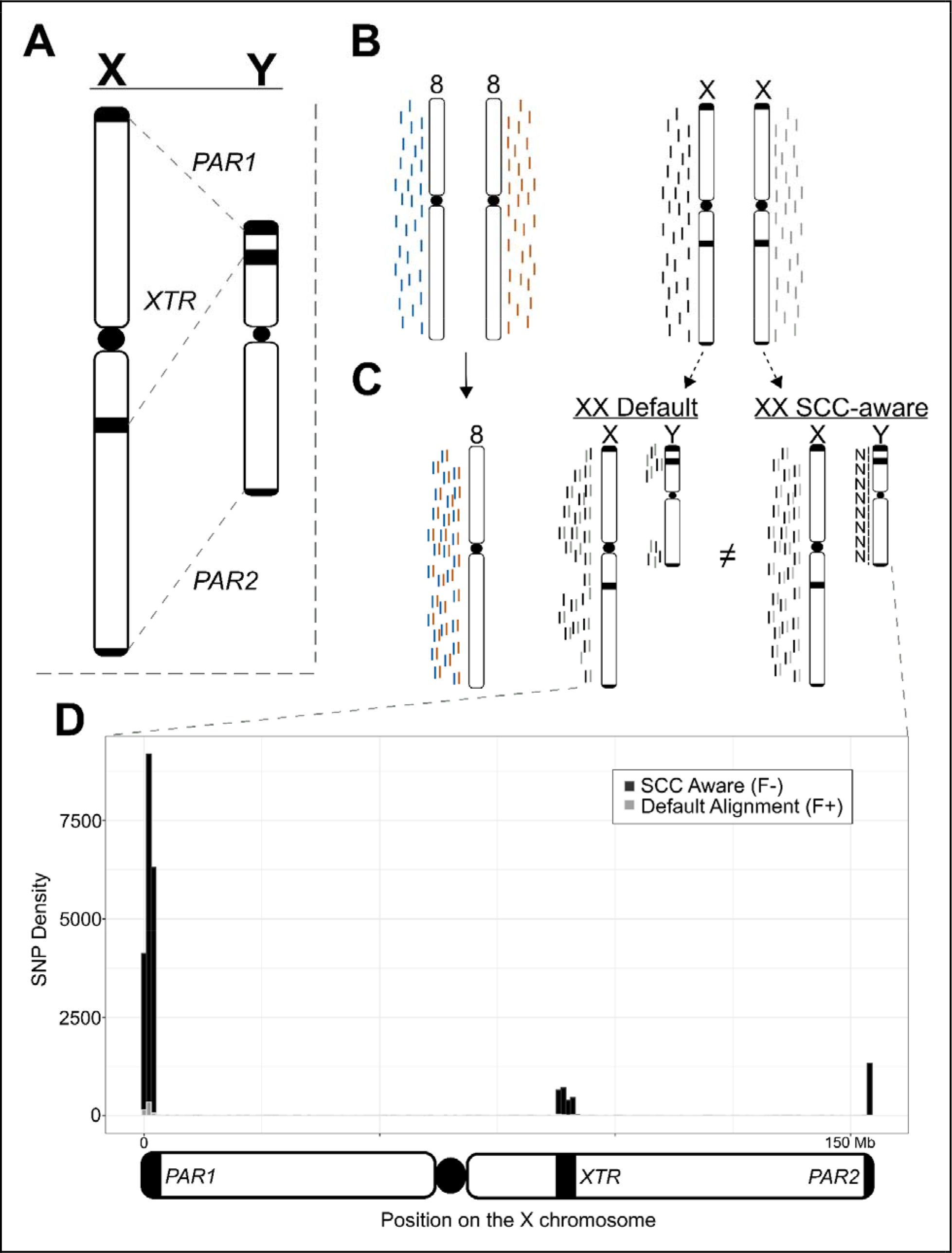
Overview of technical artifacts on the sex chromosomes for read mapping and variant calling. **(A)** Overview of regions of high sequence identity between the X and Y chromosomes. **(B)** NGS reads originating from a karyotypically diploid XX individual. **(C)** How reads from an XX-karyotype individual align to the Default reference and how masking the Y chromosome in these cases improves read mapping quality in these regions. **(D)** Connecting changes in read mapping to differences in called variants across the X chromosome in the analysis presented in this paper. Dark/black regions can be viewed as presumed as false negatives (variants missed with the Default reference), while light/gray areas can be viewed as false positive calls (variants unique to the Default reference). Overlapping variant calls between the two reference genomes have been removed. The three regions that contain the most incorrect calls using the Default reference are the two pseudoautosomal regions (beginning and end of the plot) and the X-transposed region (just right of the plot center). SNPs are binned into 1Mb windows.

The typical human genome contains a diploid count of 46 chromosomes (2n=46), but reference genome-based analyses require haploid representation of each chromosome for correct inference (e.g., n=23). In humans, the reference genome complement includes haploid representations for each autosome (n=22), but not the sex chromosomes, X and Y (n=2); thus, the human reference genome contains an n=24 chromosome representation (Figure 1b). The X and Y deviate from autosomal expectations in that (1) not all individuals possess a Y chromosome, making all reads mapping to the Y chromosome erroneous in XX (or X0 for example) samples, and (2) the X and Y retain regions of high sequence similarity (maintaining between 98-100% sequence identity) due to their shared ancestry (Olney et al. 2020; Rhie et al. 2022). Thus, particular regions on the X and Y chromosomes violate the assumption that reference genome representation for linear alignments be uniformly haploid.

According to the most recent telomere-to-telomere (T2T) human reference genome (CHMv2.0), the X chromosome makes up 5.04% of the total genome size and contains approximately the same percentage of annotated genes (Nurk et al. 2022). Thus, many published studies in humans blatantly ignore 5% or more of the human genome when conducting routine genomic analyses (Koboldt, 2020; Wise et al. 2013; Zverinova and Guryev, 2021). Indeed, despite recent advances in methodology to control for known technical artifacts inherent when analyzing the sex chromosomes (e.g., Webster et al. 2019), little progress has been made to further incorporate the X chromosome into broader biological analyses (Carey et al. 2022; Sun et al. 2023; Wise et al. 2013).

We set out to identify the extent of technical artifacts introduced by using the most complete human genome assembly currently available. Specifically, we aimed to better understand the benefits of accurately representing the sex chromosome complement when conducting standard genomic analyses. To parse the effects of the T2T-CHM13 reference genome on downstream analyses, we conducted parallel analyses using the GenBank default reference genome (Default) and a sex chromosome complement aware reference (SCC-aware) using whole genome re-sequencing and RNAseq data for 50 individuals from the Genotype-Tissue Expression (GTEx) project. We found that every analysis suffered in some capacity (in either accuracy, robustness, or both) by not using the reference genome appropriate for the data. In line with observations from previous simulation studies, we find an overwhelming number of new variants called using a SCC-aware reference that are missed when using a Default reference (Oill, 2022) that are focused in regions with higher sequence similarity to the Y chromosome than most of the X chromosome (i.e., both pseudoautosomal regions, PARs, and the X-transposed region, XTR). These differences are substantial and comprise approximately 5% of the total variants on the X chromosome.

## Methods

### Computational Overview

All primary analyses for this project were conducted on the Terra platform (Schatz et al. 2022), which interfaces multiple biomedical genomic databases with Google Cloud (GCP) through the NIH Cloud Platform Interoperability Effort (NCPI). As such, all analyses detailed below were written in Workflow Description Language (WDL); they are available for re-use here (https://github.com/DrPintoThe2nd/XYalign_AC3) and are available for integration into others’ custom Terra workspaces via Dockstore (https://dockstore.org). Further, all analyses were conducted in a single, stable Docker container (Merkel, 2014) including the following software and their dependencies (in alphabetical order): bamtools [v2.5.2] (Barnett et al. 2011), bbmap [v38.96] (Bushnell, 2014), bcftools [v1.15.1] (Li, 2011), bedtools [v2.30.0] (Quinlan and Hall, 2010), bwa [v0.7.17] (Li and Durbin, 2009), gatk4 [v4.2.6.1] (McKenna et al. 2010), hisat2 [v2.2.1] (Kim et al. 2019), openssl [v1.1.1q] (OpenSSL Project, 2003), pandas [v1.4.3] (McKinney, 2010), rtg-tools [v3.12.1] (Cleary et al. 2015), salmon [v1.9.0] (Patro et al. 2017), samblaster [v0.1.26] (Faust and Hall, 2014), samtools [v1.15.1] (Li and Durbin, 2009), Trim Galore! [v0.6.7] (Martin, 2011; https://doi.org/10.5281/zenodo.5127899). This Docker is publicly available for re-use (https://hub.docker.com/r/drpintothe2nd/ac3_xysupp).

### Data Description

We selected a subset of 50 individuals annotated as female (N=50, 46, XX) from the Genotype-Tissue Expression (GTEx) project (Aguet et al. 2020). All samples were consistent with 46,XX karyotype except for one, which we discarded due to anomalous read depth issues (adjusted N=49). Each individual possessed a minimum of whole genome re-sequencing data and RNAseq data for the same tissue; we chose the nucleus accumbens region of the basal ganglia because brain regions tend to have a high number of expressed genes (Li et al. 2017) and there is little difference in how distinct tissues are affected by reference genome mapping (Olney et al. 2020).

### Variant Calling

Because genomic data are stored on the cloud in a compressed alignment format (either CRAM or BAM—depending on data type), we first converted these files to unaligned read files, filtered PCR duplicates, and trimmed them using samtools, bbmap, and Trim Galore!, respectively. We used bwa (DNA) and hisat2 (RNA) to realign them to two different configurations of the recently published T2T human reference genome (CHM13v2.0; Nurk et al. 2022). The first configuration of the reference used was the default version downloaded from GenBank (Default), while the other was prepared as an XX-karyotype specific reference genome by hard masking the Y chromosome (Sex Chromosome Complement Aware; SCC-aware) using XYalign (Webster et al. 2019).

This type of approach also improves variant calling in XY samples (Oill, 2022; Rhie et al. 2022). As the downloaded genome version does not include a mitogenome sequence, both the Default and SCC-aware reference genomes were spiked with the mitogenome from the GRCh38 reference to help prevent mtDNA reads from mis-mapping to our regions of interest. We called variants on chromosome 8 and the X chromosome using GATK’s `HaplotypeCaller` and `GenotypeGVCFs` functions. We filtered to select only biallelic variants with: greater than or equal to four alleles present in called genotypes (`AN >= 4`), high mapping quality (`MQ > 40.0`), a minimum quality by depth of seven (`QD > 7.0`), and a total read depth of greater than or equal to ten, but less than or equal to 2500 (`DP >= 10.0 && DP <= 2500.0`). We parsed and interrogated the resultant VCF files using bcftools, rtgtools, and bedtools to better characterize the technical artifacts involved in mapping to the Default vs. SCC-aware reference genomes.

### RNAseq analyses

We analyzed the effects of reference genome on two common RNAseq data analysis, gene expression analysis and allele-specific expression analysis using salmon and GATK, respectively. We generated Default and SCC-aware reference transcriptomes for salmon analysis by extracting transcripts from the Default and SCC-aware genomes from the RefSeq annotation file using gffread [v0.12.1] (Pertea and Pertea, 2020). We soft-masked an alternate version of the Default genome using RepeatModeler [v2.0.3] (Flynn et al. 2020) to facilitate generating index decoys via the `generateDecoyTranscriptome.sh` script accompanying salmon software distribution. We ran salmon using the trimmed RNAseq reads for each individual for each reference transcriptome using the `--gcBias` and `--validateMappings` flags. For allele-specific expression (ASE), we split the filtered, genotyped VCF for each individual using bcftools and combined each individual VCF file with their re-aligned RNAseq data using GATK’s `ASEReadCounter` function. We compared results between reference genomes as a deviation from a 1:1 relationship. For ASE, we also compared the efficacy of variant calling and alignment on the total number of transcripts identified as allele-specific.

## Results

### Sex chromosome aware reference augments variant calling

At a broad scale, we identified that the SCC-aware reference alignment increased the number of properly paired reads mapped for many individuals (mean: +6,551; +0.0008%) and decreased in mapped reads with a mapping quality of 0 (`MQ=0`) in every individual (mean: −605,396; −1.05%) (Supplemental Tables 1 and 2). These changes in read mapping resulted in changes in the total number of biallelic, single nucleotide variants (SNPs) among all 49 individuals. In contrast, on chromosome 8 the total number of variants called were nearly identical between the two reference genome configurations—719,826 variants and 719,824 variants for the Default and SCC-aware reference, respectively. At a per-individual scale, this course held with average numbers of variants being 178,885 and 178,882 variants, respectively (Figure 1; Table 1).

**Table 1:** Numerical differences in variant calling outcomes on chromosome 8 and the X chromosome between sex chromosome complement-aware (SCC-aware) and default reference alignment. Numbers are quality-filtered biallelic SNPs for chromosome 8 (top) and the X chromosome (bottom).

| Category | Chrom | Default | SCC-aware | % change (SCC/D) |
| --- | --- | --- | --- | --- |
| Total SNPs | 8 | 719,826 | 719,824 | -0.0002% |
| Per-indiv. Avg SNPs | 8 | 178,885 | 178,882 | -0.002% |
| Per-indiv. Ref allele | 8 | 540,938 | 540,939 | 0.0002% |
| Total SNPs | X | 475,763 | 498,297 | 4.74% |
| Per-indiv. Avg SNPs | X | 98,877 | 105,413 | 6.61% |
| Per-indiv. Ref allele | X | 376,884 | 392,882 | 4.07% |

However, this impartiality was not replicated on the X chromosome, where we found a sharp increase of 22,534 total SNPs (from 475,763 to 498,297) when using the SCC-aware reference configuration. This deviation also held for each individual in our study, with an average increase in the number of called SNPs from 98,877 to 105,413 (Table 1; Supplemental Table 3).

Across most of the X chromosome (~95%), we found little variation between the two reference genome configurations (Figure 1d; Table 2). Indeed, as most of the X chromosome shares little sequence identity between the X and Y chromosomes, very few areas generate read mapping conflict between them, even for XX samples (Figure 1). In the three regions of high sequence similarity (PAR1, XTR, and PAR2), changes in total numbers of SNPs called between reference configurations ranged from an 11% increase to a 730% increase in the XTR and PARs, respectively (Table 2). Indeed, we saw an increase in called variants in both genic (PARs: +564.39%; XTR: +13.59%) and intergenic (PARs: +894.71%; XTR: +10.37) regions (Table 2). Thus, while differences across most of the X chromosome are negligible, the differences in numbers of called SNPs in the XTR and two PARs are significant relative to both the autosomes or the rest of the X chromosome.

**Table 2:** Dissection of differences in variant calling within regions of interest across the X chromosome. Numbers are quality-filtered biallelic SNPs across various regions on the X chromosome (top) and within genic regions only across various regions on the X chromosome (bottom).

| Category | Default | SCC-aware | % change (SCC/D) | Added (F-) | Lost (F+) |
| --- | --- | --- | --- | --- | --- |
| **Total SNPs** | 475,763 | 498,297 | +4.74% | 23,279 | 745 |
| non-PAR/XTR | 453,822 | 453,882 | +0.01% | 150 | 90 |
| PARs | 2,790 | 23,161 | +730.14% | 20,931 | 560 |
| XTR | 19,151 | 21,254 | +10.98% | 2,198 | 95 |
| **Genic SNPs** | 162,989 | 171,351 | +5.13% | 8,600 | 238 |
| non-PAR/XTR | 157,957 | 157,979 | +0.01% | 45 | 23 |
| PARs | 1,390 | 9,235 | +564.39% | 8,042 | 197 |
| XTR | 3,642 | 4,137 | +13.59% | 513 | 18 |

### Default reference distorts gene expression quantification on the X

Somewhat contrary to the exceptional differences between variant calling with different reference genome configurations, differences between gene expression quantification are more subtle, yet still apparent (Figure 2). For gene expression quantification, we calculated Transcripts Per Kilobase Million (TPM) and found that differences between expression levels are greatest in PAR1, followed by PAR2, and then the rest of the chromosome (Supplemental Figure 3a). However, contrary to expectations we find little changes in expression values within the XTR (Supplemental Figure 3a). Also contrary to expectations, we find no relationship between observed expression differences and transcript length (Supplemental Figure 3b) or expression level (Supplemental Figure 3c-d).

**Figure 2:**
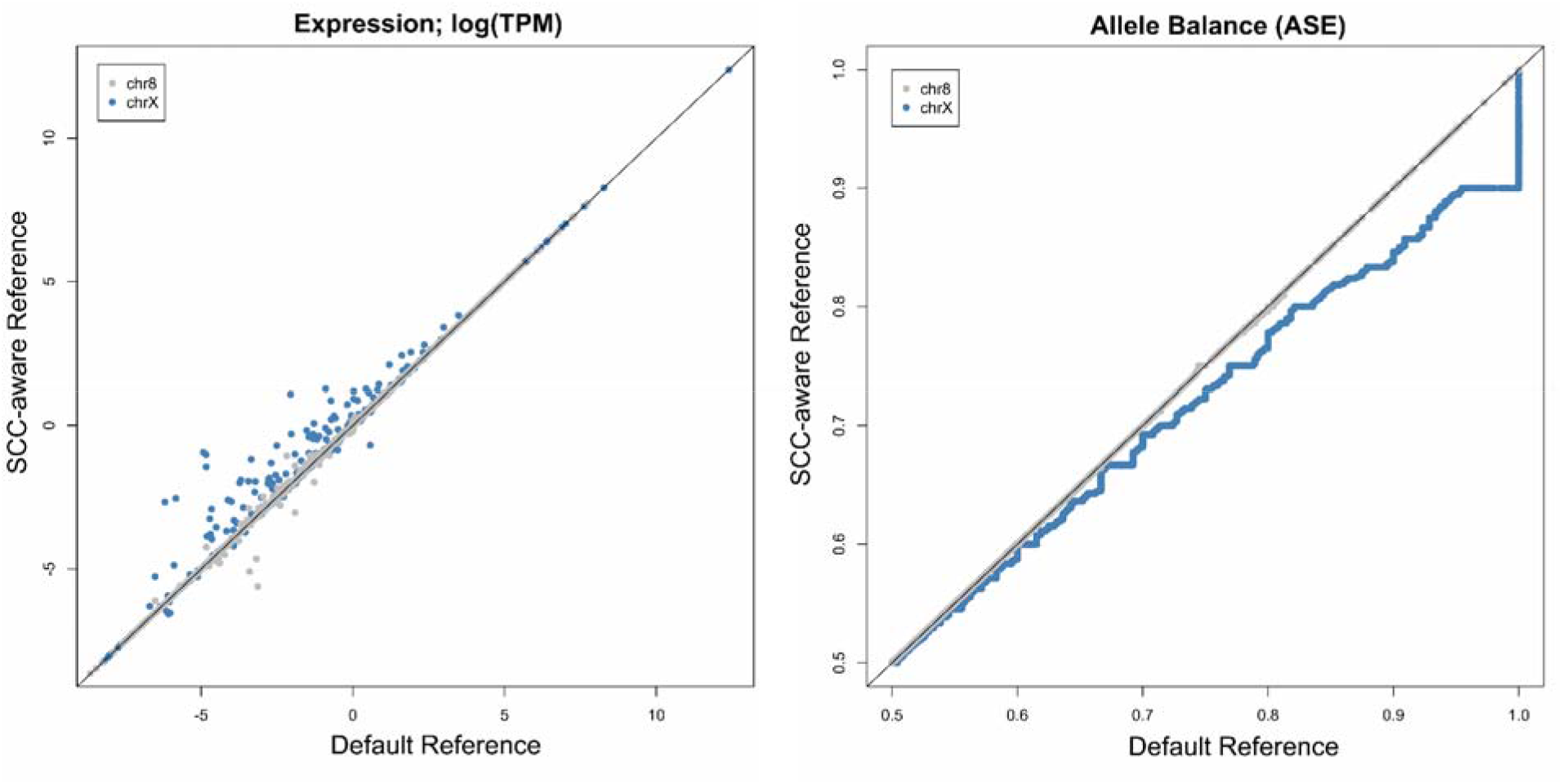
Effects of the SCC-aware reference genome on common RNAseq analyses: (left) gene expression (Transcripts Per Kilobase Million; TPM) and (right) allele balance (allele-specific expression; ASE). For allele balance, we used the SCC-aware reference called VCF as a measure to increase the total number of transcripts included (see Table 3). Both analyses use the T2T-CHM13v2 genome sequence for mapping.

When examining allele-specific expression (ASE) levels, or the allele-balance ratio, we see an opposite pattern—where the higher expressed a transcript is, the more skewed the Default alignment data become on the X chromosome. We observed that allele balance values are generally inflated using the Default reference (Figure 2). Importantly, we see a premature summit, or abbreviated climb, from allele balance values from 0.5 to 1.0, when using the Default reference genome—where allele balance values >0.9 get rounded up to 1.0 (Figure 2). Because there is an extra alignment step in ASE analysis relative to regular expression quantification (i.e., variant calling), we attempted to parse which aspects of ASE analysis are most affected by which segment of the analysis. We paired each potential variant calling output (VCF file) with each potential RNAseq alignment output (BAM file) by re-running the analysis in a “round-robin”, or “all-vs-all”, format. We found that the VCF file (and thus the reference genome used for variant calling) chosen to run ASE had the greatest influence on the number recovered biallelic transcripts (Table 3).

**Table 3:** Efficacy of allele-specific expression (ASE) analysis across differing modes of variant calling and RNAseq alignment strategies.

| ASE Mode | chr8 # | chrX # |
| --- | --- | --- |
| Default VCF, Default RNAseq | 885 | 819 |
| Default VCF, SCC-aware RNAseq | 885 | 823 |
| SCC-aware VCF, Default RNAseq | 885 | 834 |
| SCC-aware VCF, SCC-aware RNAseq | 885 | 835 |

## Discussion

As expected, there were negligible differences in all analyses between results on chromosome 8 between Default and SCC-aware reference genomes (Figure 2; Tables 1 & 3). However, the differences on the X chromosome were substantial (Figure 1–2; Tables 1–3). The most numerous differences between the Default and SCC-aware reference genomes were the sheer number of (presumed) false negatives when using the Default reference, i.e., variants called using the SCC-aware reference but missed with the Default reference (Table 2). There were also (presumed) false positives, variants called with the Default reference that were absent in the SCC-aware reference; however, these made up a small fraction of the observed differences (Table 2). To expand on this concept, we calculated the major allele frequencies for all sites in both the Default and SCC-aware VCFs (Supplemental Figure 1) and then filtered out variants that overlap between the two (Supplemental Figure 2). We expected that if one spectrum contained an increase in false positives the major allele frequency would skew more heavily towards 1.0 (an increase in singleton calls). Indeed, this is exactly what we observed in both PAR regions and the XTR (Supplemental Figures 1 and 2).

Although the pseudoautosomal regions (PARs) make-up only ~1.77% of the X chromosome, they contain ~5% of both genic (5.39%) and indiscriminate (all) SNPs (4.65%) within our sampled individuals. However, using the Default reference genome, these numbers are unfathomably low for both genic (0.85%) and indiscriminate SNPs (0.59%). This pattern also holds, albeit mediated by genetic divergence between X and Y alleles relative to the PARs, within the X transposed region (XTR). The XTR makes up ~3.04% of the X chromosome, yet the numbers of called SNPs increase substantially when using the appropriate SCC-aware reference compared to the Default for both genic (2.2% to 2.4%) and indiscriminate SNPs (4.0% to 4.3%).

Our expression analyses of RNAseq data may be the first published RNAseq analyses using the CHM13_v2.0 assembly. Our comparative expression analysis suggests that a notable amount of gene expression differences can be found throughout the X chromosome but are most notable in PAR1 (Supplemental Figure 3a). Interestingly, we note that allele-specific expression (ASE) analysis especially suffers from a two-fold increase in error when using an inappropriate reference genome. The first introduction of error, as mentioned previously, is the substantial number of false negatives introduced during variant calling via mapping WGS reads (Tables 1 & 2). The second error is introduced during mapping RNAseq reads to the Default reference, whereby correcting for either factor (called SNPs or RNAseq mapping) can partially recover some of the potentially missed transcripts in an ASE experiment (Table 3). However, to take full advantage of ASE analyses on the X chromosome, it is essential to include both correctly called variants and correctly mapped RNAseq reads (Table 3; Supplemental Table 4).

In line with previous conclusions (e.g., Wise et al. 2013), the general absence of the X chromosome in many analyses may be due, in part, to an increase in technical effort/ability to prepare the reference genome prior to analysis (Webster et al. 2019). The X chromosome makes up 5% of the haploid genome size of the typical XX human individual. Therefore, the “scorched earth” error rate of not including the X chromosome in genomics analyses of XX individuals is at least 5%. The introduction of read mapping errors on the X chromosome only affects 5% of the total length of the X chromosome, which equates to only 0.25% of the variants called become unreliable when not accounting for sex chromosome complement and using a Default reference genome (Figure 1; Table 2). Thus, the common practice of purposefully introducing an error rate of 5% (excluding the X chromosome) to potentially avoid an error rate of 0.25% (including the X chromosome) is excessive and, technically speaking, precludes the use of the term “genome-wide” in most association studies in humans (Sun et al. 2023; Wise et al. 2013). However, it is a relatively trivial task to inform the reference genome with the sex chromosome complement when mapping samples and accommodate changes in ploidy across different regions; thus ensuring that reliable variant calls across, even within the PARs and XTR (Carey et al. 2022; Webster et al. 2019). We expect the broader utilization of the SCC-aware reference genome for alignment could be catalyzed by it being made available alongside the Default on repositories such as NCBI’s GenBank, where the main hurdle to its inclusion may be low (Carey et al. 2022).

In conclusion, we conducted a pilot study of replicating a series of commonly used genomics tools/analyses across a subset of the GTEx data available on the cloud. We showed that technical artifacts introduced by using the Default reference genome affect about 5% across the X chromosome but are most extensive in the PARs and XTR, ranging upwards of 700% in some regions. In line with prior work, we provided additional evidence that technical artifacts of including the sex chromosomes in genomics analyses can be negated with available information and tools (Olney et al. 2019; Webster et al. 2019). We know that, though the ‘eXclusion’ of the X chromosome is widespread (Wise et al. 2013), the exclusion of the Y is even more extensive in empirical and clinical genomics (Sun et al. 2023). SCC-aware reference genomes can effectively negate the effects of homology on the sex chromosomes in XX individuals and reduce this mis-mapping in XY individuals, allowing for their accurate inclusion in human genomics studies (Oill, 2022). We’re hopeful that research groups will make the inclusion of SCC-aware references a staple in their future projects—not only to better reflect the original intent behind the National Institutes of Health of the USA’s policy on the consideration of sex as a biological variable (https://orwh.od.nih.gov/sex-gender/nih-policy-sex-biological-variable), but also to bring humanity a better understanding of how sex chromosomes affect human health and disease states across the world.

## Acknowledgements

We acknowledge Research Computing at Arizona State University for providing high-performance computing and storage resources that have contributed to the research results reported within this paper (http://www.researchcomputing.asu.edu). We would also like to thank Wilson lab members for helpful feedback on the research and manuscript. This work was supported by the National Institute of General Medical Sciences (NIGMS) of the National Institutes of Health grant R35GM124827 to M.A.W. Funding for the computational aspects of this research was provided by the AnVIL Cloud Credits (AC3) pilot program for computation using the Terra platform to M.A.W.

## Data Availability

The data used in this study are available as follows, reference genome T2T-CHM13v2.0, GenBank: GCA_009914755.4;The Genotype-Tissue Expression (GTEx) Project was supported by the Common Fund of the Office of the Director of the National Institutes of Health, and by NCI, NHGRI, NHLBI, NIDA, NIMH, and NINDS. The GTEx data is described and available through dbGaP under accession phs000424.v8.p1; we received approval to access this data under dbGaP accession #8834; and code to replicate results on the Terra cloud computing environment on GitHub/Dockstore (https://github.com/DrPintoThe2nd/XYalign_AC3).

