## Supplemental for "Concerning the eXclusion in human genomics: The choice of sex chromosome representation in the human genome drastically affects number of identified variants"

### Supplementary Materials

**Supplemental Figure 1:** Major allele frequency plots for all SNPs in each of the three X chromosome regions that possess high sequence similarity to the Y chromosome.

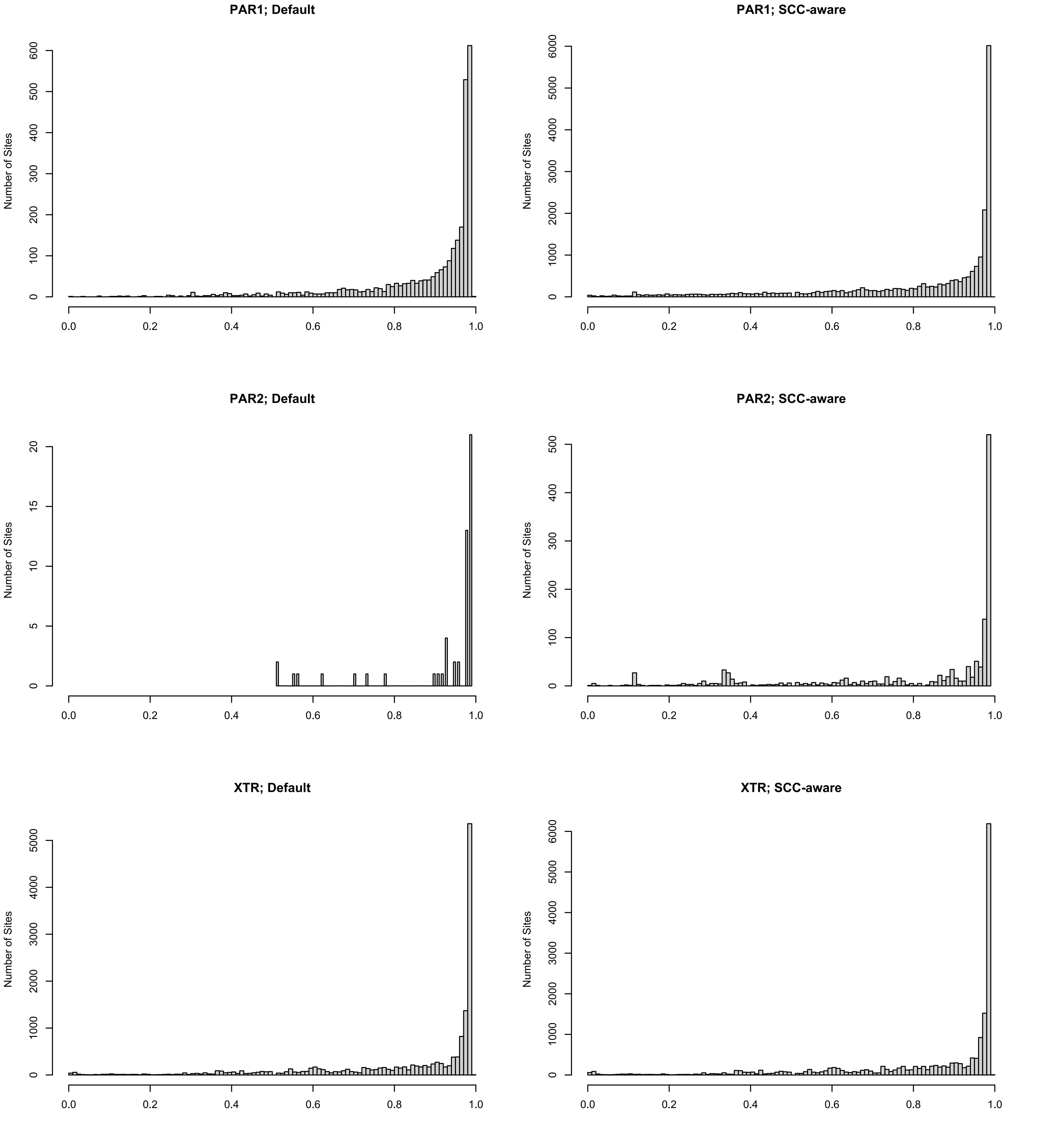
PAR1; Default
PAR1; SCC-aware
PAR2; Default
PAR2; SCC-aware
XTR; Default
XTR; SCC-aware
Number of Sites
0.0
0.2
0.4
0.6
0.8
1.0

**Supplemental Figure 2:** Major allele frequency plots for non-overlapping SNPs between the two reference configurations in each of the three X chromosome regions that possess high sequence similarity to the Y chromosome.

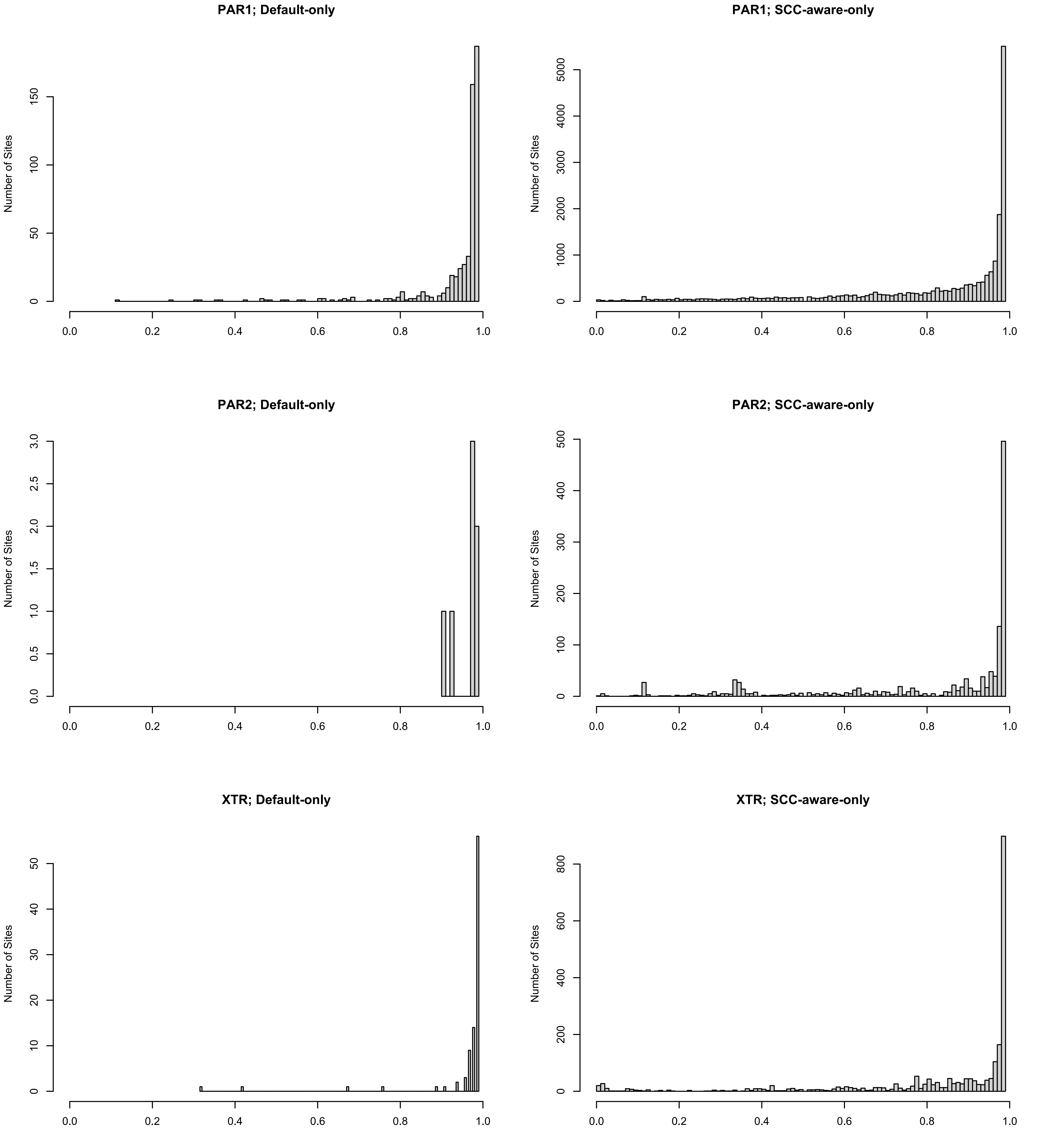

PAR1; Default-only
PAR1; SCC-aware-only
PAR2; Default-only
PAR2; SCC-aware-only
XTR; Default-only
XTR; SCC-aware-only
Number of Sites
0.0
0.2
0.4
0.6
0.8
1.0

**Supplemental Figure 3:** Interrogating expression differences across the X chromosome. **(A)** We find that false changes in expression values are (i) nearly absent in the XTR, (ii) greatest in the PARs, and (iii) occur sporadically throughout regions with little sequence similarity with the Y. **(B)** No relationship between total transcript length and differences in expression values. There was also no observed relationship between expression levels and observed expression differences with either the **(C)** Default or **(D)** SCC-aware reference transcriptomes.

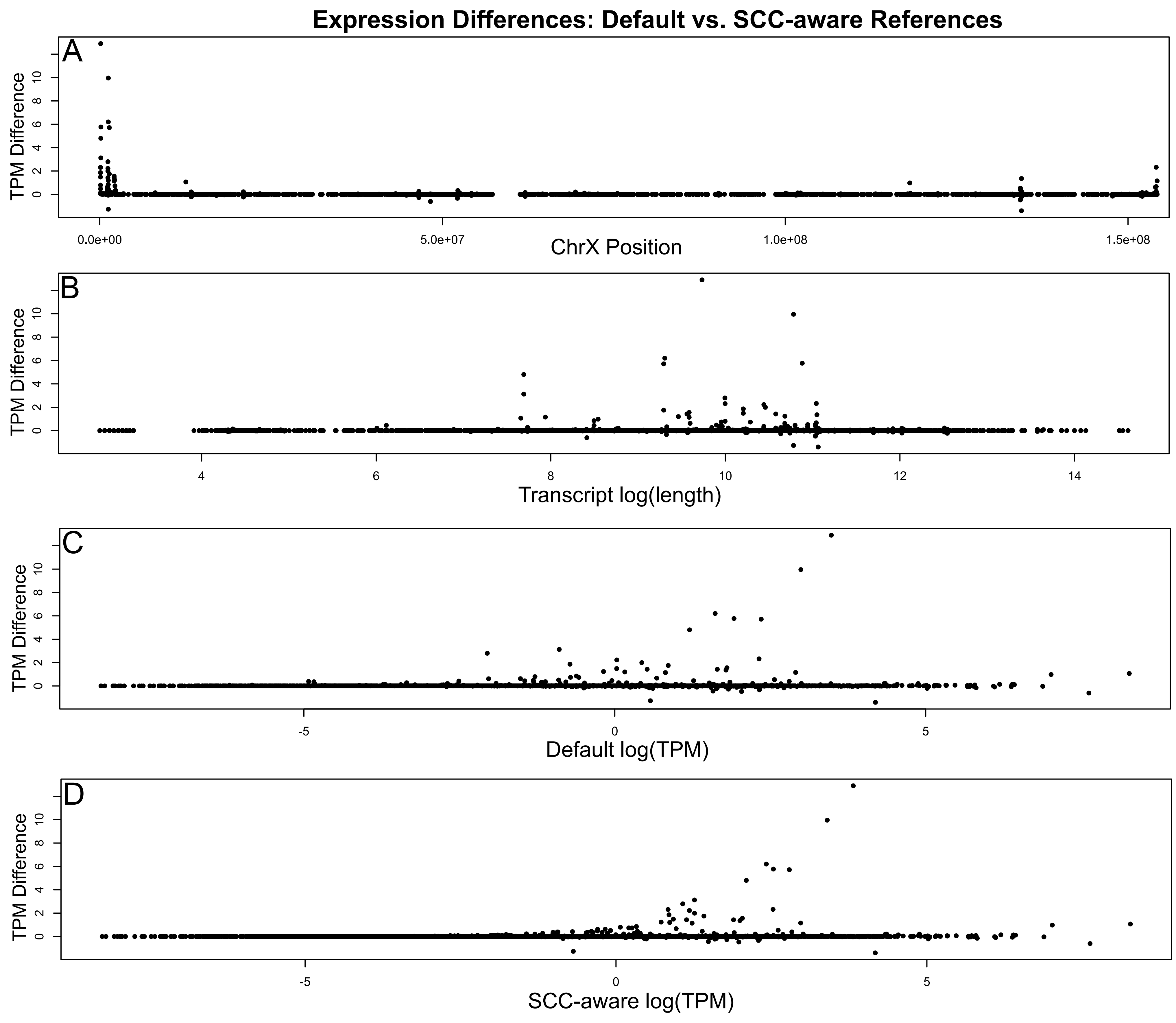

Expression Differences: Default vs. SCC-aware References
A
TPM Difference
ChrX Position
0.0e+00
5.0e+07
1.0e+08
1.5e+08
B
Transcript log(length)
C
Default log(TPM)
D
SCC-aware log(TPM)

**<u>Supplemental Table 1</u>:** Numbers of properly paired reads aligned per-individual and their differences (Diff == SCC-aware - Default).

| Sample | Default | SCC-aware | Diff |
|---|---|---|---|
| GTEX-11GSP | 743,538,052 | 743,514,590 | -23462 |
| GTEX-11TTK | 762,276,120 | 762,263,034 | -13086 |
| GTEX-12WSD | 779,860,512 | 779,877,012 | 16500 |
| GTEX-12ZZX | 770,109,140 | 770,094,676 | -14464 |
| GTEX-1313W | 852,501,364 | 852,498,722 | -2642 |
| GTEX-131XW | 718,436,724 | 718,428,810 | -7914 |
| GTEX-13JUV | 850,778,578 | 850,773,524 | -5054 |
| GTEX-13O3O | 678,903,264 | 679,008,546 | 105282 |
| GTEX-13OVJ | 657,151,114 | 657,202,072 | 50958 |
| GTEX-13PL6 | 807,524,294 | 807,517,724 | -6570 |
| GTEX-13QJC | 752,669,274 | 752,663,422 | -5852 |
| GTEX-13S7M | 871,498,252 | 871,491,502 | -6750 |
| GTEX-13SLX | 713,876,242 | 714,000,110 | 123868 |
| GTEX-13X6K | 829,626,906 | 829,620,008 | -6898 |
| GTEX-145MI | 747,642,106 | 747,638,062 | -4044 |

| | | | |
|---|---|---|---|
| GTEX-14BIM | 807,883,826 | 807,878,368 | -5458 |
| GTEX-14BIN | 746,761,356 | 746,748,214 | -13142 |
| GTEX-14LZ3 | 814,097,488 | 814,090,184 | -7304 |
| GTEX-14PJM | 636,542,840 | 636,587,472 | 44632 |
| GTEX-14PQA | 793,581,622 | 793,576,584 | -5038 |
| GTEX-15DCD | 785,001,666 | 784,994,964 | -6702 |
| GTEX-15DYW | 769,319,014 | 769,312,500 | -6514 |
| GTEX-15ER7 | 1,032,636,384 | 1,032,596,560 | -39824 |
| GTEX-17JCI | 722,476,160 | 722,468,042 | -8118 |
| GTEX-183WM | 644,882,656 | 644,859,502 | -23154 |
| GTEX-1B933 | 786,036,470 | 786,029,174 | -7296 |
| GTEX-1CB4H | 1,023,380,652 | 1,023,367,536 | -13116 |
| GTEX-1EWIQ | 803,716,524 | 803,712,240 | -4284 |
| GTEX-1F48J | 773,146,722 | 773,139,548 | -7174 |
| GTEX-1F7RK | 677,735,730 | 677,934,982 | 199252 |
| GTEX-1F88E | 769,156,638 | 769,153,806 | -2832 |
| GTEX-1GMR8 | 736,925,836 | 736,922,506 | -3330 |
| GTEX-1GN1U | 724,367,780 | 724,358,848 | -8932 |

| | | | |
|---|---|---|---|
| GTEX-1GZHY | 777,817,298 | 777,808,354 | -8944 |
| GTEX-1LH75 | 677,035,540 | 677,033,288 | -2252 |
| GTEX-1OJC4 | 718,447,988 | 718,439,508 | -8480 |
| GTEX-1R46S | 729,160,502 | 729,151,946 | -8556 |
| GTEX-N7MT | 1,101,259,154 | 1,101,389,954 | 130800 |
| GTEX-NPJ7 | 1,189,397,988 | 1,189,402,522 | 4534 |
| GTEX-PWO3 | 1,505,297,004 | 1,505,284,312 | -12692 |
| GTEX-Q2AG | 1,460,521,486 | 1,460,507,080 | -14406 |
| GTEX-QDT8 | 734,468,868 | 734,467,724 | -1144 |
| GTEX-QVJO | 936,001,950 | 935,995,918 | -6032 |
| GTEX-QVUS | 1,022,722,188 | 1,022,716,138 | -6050 |
| GTEX-RU72 | 951,784,008 | 951,778,084 | -5924 |
| GTEX-T2IS | 894,646,420 | 894,641,480 | -4940 |
| GTEX-TSE9 | 882,487,064 | 882,481,826 | -5238 |
| GTEX-WWYW | 1,004,805,488 | 1,004,799,550 | -5938 |
| GTEX-X4EP | 940,866,246 | 940,860,976 | -5270 |
| **Average:** | **838,995,112** | **839,001,663** | **6551** |

**Supplemental Table 2:** Numbers of aligned reads with a mapping quality score of 0 per-individual and their differences (Diff == SCC-aware - Default).

| Sample | Default | SCC-aware | Diff |
|---|---|---|---|
| GTEX-11GSP | 52,587,267 | 51,963,148 | -624119 |
| GTEX-11TTK | 57,280,032 | 56,705,681 | -574351 |
| GTEX-12WSD | 63,284,582 | 62,699,648 | -584934 |
| GTEX-12ZZX | 63,128,131 | 62,555,073 | -573058 |
| GTEX-1313W | 56,811,303 | 56,257,030 | -554273 |
| GTEX-131XW | 49,082,779 | 48,572,485 | -510294 |
| GTEX-13JUV | 58,887,315 | 58,332,639 | -554676 |
| GTEX-13O3O | 59,712,155 | 59,168,081 | -544074 |
| GTEX-13OVJ | 52,722,912 | 52,207,406 | -515506 |
| GTEX-13PL6 | 58,661,009 | 58,119,562 | -541447 |
| GTEX-13QJC | 54,114,368 | 53,593,977 | -520391 |
| GTEX-13S7M | 54,699,099 | 54,117,601 | -581498 |
| GTEX-13SLX | 59,019,631 | 58,446,241 | -573390 |
| GTEX-13X6K | 56,917,952 | 56,386,489 | -531463 |
| GTEX-145MI | 52,799,420 | 52,298,758 | -500662 |

| | | | |
|---|---|---|---|
| GTEX-14BIM | 61,610,959 | 61,030,765 | -580194 |
| GTEX-14BIN | 43,207,035 | 42,559,653 | -647382 |
| GTEX-14LZ3 | 62,991,712 | 62,403,932 | -587780 |
| GTEX-14PJM | 49,534,691 | 48,982,854 | -551837 |
| GTEX-14PQA | 50,100,257 | 49,540,704 | -559553 |
| GTEX-15DCD | 47,669,993 | 47,063,323 | -606670 |
| GTEX-15DYW | 43,796,294 | 43,232,473 | -563821 |
| GTEX-15ER7 | 74,279,573 | 73,529,304 | -750269 |
| GTEX-17JCI | 68,252,280 | 67,695,300 | -556980 |
| GTEX-183WM | 50,108,997 | 49,655,847 | -453150 |
| GTEX-1B933 | 54,902,828 | 54,367,966 | -534862 |
| GTEX-1CB4H | 75,562,943 | 74,817,585 | -745358 |
| GTEX-1EWIQ | 55,647,825 | 55,080,569 | -567256 |
| GTEX-1F48J | 56,413,907 | 55,844,076 | -569831 |
| GTEX-1F7RK | 56,474,139 | 55,910,411 | -563728 |
| GTEX-1F88E | 28,710,651 | 28,338,932 | -371719 |
| GTEX-1GMR8 | 51,148,341 | 50,661,769 | -486572 |
| GTEX-1GN1U | 55,599,531 | 55,075,668 | -523863 |

| | | | |
|---|---|---|---|
| GTEX-1GZHY | 58,201,555 | 57,655,445 | -546110 |
| GTEX-1LH75 | 33,992,212 | 33,621,491 | -370721 |
| GTEX-1OJC4 | 40,547,630 | 39,981,337 | -566293 |
| GTEX-1R46S | 44,843,832 | 44,371,247 | -472585 |
| GTEX-N7MT | 82,630,657 | 81,661,679 | -968978 |
| GTEX-NPJ7 | 89,262,358 | 88,282,682 | -979676 |
| GTEX-PWO3 | 100,949,535 | 99,828,089 | -1121446 |
| GTEX-Q2AG | 99,396,799 | 98,307,124 | -1089675 |
| GTEX-QDT8 | 24,714,409 | 24,395,809 | -318600 |
| GTEX-QVJO | 56,956,907 | 56,279,207 | -677700 |
| GTEX-QVUS | 67,919,175 | 67,198,696 | -720479 |
| GTEX-RU72 | 67,284,764 | 66,662,729 | -622035 |
| GTEX-T2IS | 52,908,632 | 52,188,901 | -719731 |
| GTEX-TSE9 | 58,283,906 | 57,678,080 | -605826 |
| GTEX-WWYW | 65,204,859 | 64,483,930 | -720929 |
| GTEX-X4EP | 57,656,597 | 56,997,932 | -658665 |
| **Average:** | **57,887,219** | **57,281,823** | **-605396** |

**Supplemental Table 3:** Per individual number of SNPs called noting no outlier individuals suggesting potentially false positives. Diff=SCC[-aware]-Def[ault]

| Sample | chr8 (Def) | chr8 (SCC) | chr8 (Diff) | chrX (Def) | chrX (SCC) | chrX (Diff) |
|---|---|---|---|---|---|---|
| GTEX-11GSP | 170326 | 170324 | -2 | 93320 | 99972 | 6652 |
| GTEX-11TTK | 175931 | 175921 | -10 | 88627 | 95224 | 6597 |
| GTEX-12WSD | 226710 | 226708 | -2 | 148019 | 156376 | 8357 |
| GTEX-12ZZX | 178513 | 178513 | 0 | 94381 | 100696 | 6315 |
| GTEX-1313W | 174948 | 174945 | -3 | 93046 | 99266 | 6220 |
| GTEX-131XW | 177765 | 177766 | 1 | 92975 | 99458 | 6483 |
| GTEX-13JUV | 237761 | 237761 | 0 | 156218 | 164340 | 8122 |
| GTEX-13O3O | 177356 | 177337 | -19 | 97231 | 103865 | 6634 |
| GTEX-13OVJ | 176250 | 176242 | -8 | 92377 | 98675 | 6298 |
| GTEX-13PL6 | 184386 | 184384 | -2 | 99694 | 106357 | 6663 |
| GTEX-13QJC | 174202 | 174198 | -4 | 94956 | 101369 | 6413 |
| GTEX-13S7M | 176781 | 176773 | -8 | 97856 | 104586 | 6730 |
| GTEX-13SLX | 227689 | 227679 | -10 | 158971 | 167289 | 8318 |
| GTEX-13X6K | 176829 | 176833 | 4 | 98562 | 105018 | 6456 |

| | | | | | | |
|---|---|---|---|---|---|---|
| GTEX-145MI | 180230 | 180221 | -9 | 96266 | 102756 | 6490 |
| GTEX-14BIM | 174321 | 174318 | -3 | 94983 | 101611 | 6628 |
| GTEX-14BIN | 175646 | 175637 | -9 | 92269 | 98852 | 6583 |
| GTEX-14LZ3 | 168114 | 168112 | -2 | 97801 | 104341 | 6540 |
| GTEX-14PJM | 179596 | 179590 | -6 | 92756 | 99146 | 6390 |
| GTEX-14PQA | 174755 | 174754 | -1 | 96647 | 103237 | 6590 |
| GTEX-15DCD | 176549 | 176548 | -1 | 97316 | 103664 | 6348 |
| GTEX-15DYW | 174323 | 174313 | -10 | 92837 | 99687 | 6850 |
| GTEX-15ER7 | 176037 | 176024 | -13 | 95542 | 101525 | 5983 |
| GTEX-17JCI | 172198 | 172194 | -4 | 89354 | 95742 | 6388 |
| GTEX-183WM | 173002 | 173002 | 0 | 99420 | 105737 | 6317 |
| GTEX-1B933 | 170274 | 170270 | -4 | 98125 | 104067 | 5942 |
| GTEX-1CB4H | 177861 | 177864 | 3 | 89546 | 95366 | 5820 |
| GTEX-1EWIQ | 175684 | 175685 | 1 | 97071 | 103582 | 6511 |
| GTEX-1F48J | 175759 | 175758 | -1 | 95263 | 101619 | 6356 |
| GTEX-1F7RK | 174431 | 174430 | -1 | 100467 | 107217 | 6750 |
| GTEX-1F88E | 176955 | 176947 | -8 | 99597 | 105950 | 6353 |
| GTEX-1GMR8 | 174315 | 174301 | -14 | 93455 | 100245 | 6790 |

| | | | | | | |
|---|---|---|---|---|---|---|
| GTEX-1GN1U | 180183 | 180174 | -9 | 101162 | 107631 | 6469 |
| GTEX-1GZHY | 176848 | 176844 | -4 | 94084 | 100236 | 6152 |
| GTEX-1LH75 | 177424 | 177426 | 2 | 96356 | 102311 | 5955 |
| GTEX-1OJC4 | 173138 | 173136 | -2 | 94733 | 101247 | 6514 |
| GTEX-1R46S | 175986 | 175985 | -1 | 91872 | 98609 | 6737 |
| GTEX-N7MT | 173229 | 173226 | -3 | 94344 | 101030 | 6686 |
| GTEX-NPJ7 | 174748 | 174756 | 8 | 96675 | 102994 | 6319 |
| GTEX-PWO3 | 168746 | 168746 | 0 | 92403 | 98820 | 6417 |
| GTEX-Q2AG | 176197 | 176193 | -4 | 95794 | 102046 | 6252 |
| GTEX-QDT8 | 178424 | 178426 | 2 | 98435 | 104887 | 6452 |
| GTEX-QVJO | 167814 | 167813 | -1 | 89697 | 95740 | 6043 |
| GTEX-QVUS | 177249 | 177249 | 0 | 93340 | 99673 | 6333 |
| GTEX-RU72 | 179638 | 179638 | 0 | 98649 | 105135 | 6486 |
| GTEX-T2IS | 177434 | 177434 | 0 | 96054 | 102236 | 6182 |
| GTEX-TSE9 | 172480 | 172478 | -2 | 97895 | 104231 | 6336 |
| GTEX-WWYW | 178286 | 178293 | 7 | 94301 | 100727 | 6426 |
| GTEX-X4EP | 172029 | 172029 | 0 | 94240 | 100860 | 6620 |
| **Average:** | **178884.7** | **178881.6** | **-3.1** | **98877.2** | **105413.2** | **6536.0** |

**Supplemental Table 4**: List of ASE transcript dropouts not recovered using the Default genome configuration but recovered using the SCC-aware reference alignment.

| Chr | Start | Stop | Transcript | Source | Type | Gene |
|---|---|---|---|---|---|---|
| X | 104530 | 131585 | rna-NM_001370370.1 | BestRefSeq | mRNA | PLCXD1 |
| X | 109615 | 131585 | rna-NM_018390.4 | BestRefSeq | mRNA | PLCXD1 |
| X | 1116125 | 1155344 | rna-NM_001161531.2 | BestRefSeq | mRNA | CSF2RA |
| X | 1116125 | 1159654 | rna-NM_001161529.2 | BestRefSeq | mRNA | CSF2RA |
| X | 1116125 | 1172713 | rna-NM_001379161.1 | BestRefSeq | mRNA | CSF2RA |
| X | 1116125 | 1174802 | rna-NM_001379159.1 | BestRefSeq | mRNA | CSF2RA |
| X | 111712 | 131585 | rna-NM_001370373.1 | BestRefSeq | mRNA | PLCXD1 |
| X | 1229797 | 1235760 | rna-NM_001636.4 | BestRefSeq | mRNA | SLC25A6 |
| X | 1393021 | 1403889 | rna-NM_005088.3 | BestRefSeq | mRNA | AKAP17A |
| X | 1396844 | 1441907 | rna-NM_004043.3 | BestRefSeq | mRNA | ASMT |
| X | 1.54E+08 | 1.54E+08 | rna-NM_001394353.1 | BestRefSeq | mRNA | SPRY3 |
| X | 1.54E+08 | 1.54E+08 | rna-NM_005840.4 | BestRefSeq | mRNA | SPRY3 |
| X | 1.54E+08 | 1.54E+08 | rna-NM_001145149.3 | BestRefSeq | mRNA | VAMP7 |
| X | 1.54E+08 | 1.54E+08 | rna-XM_047447120.1 | Gnomon | mRNA | LOC124908551 |

| | | | | | | |
|---|---|---|---|---|---|---|
| X | 2304267 | 2349424 | rna-NM_001321367.2 | BestRefSeq | mRNA | CD99 |
| X | 2304267 | 2354297 | rna-NM_001122898.3 | BestRefSeq | mRNA | CD99 |
